# VHL inactivation without hypoxia is sufficient to achieve genome hypermethylation

**DOI:** 10.1101/093310

**Authors:** Artem V. Artemov, Nadezhda Zhigalova, Svetlana Zhenilo, Alexander M. Mazur, Egor B. Prokhortchouk

## Abstract

VHL inactivation is a key oncogenic event for renal carcinomas. In normoxia, VHL suppresses HIF1a-mediated response to hypoxia. It has previously been shown that hypoxic conditions inhibit TET-dependent hydroxymethylation of cytosines and cause DNA hypermethylation at gene promoters. In this work, we performed VHL inactivation by CRISPR/Cas9 and studied its effects on gene expression and DNA methylation. We showed that even without hypoxia, VHL inactivation leads to hypermethylation of the genome which mainly occurred in AP-1 and TRIM28 binding sites. We also observed promoter hypermethylation of several transcription regulators associated with decreased gene expression.

## Introduction

Sequencing of cancer genomes was initially aimed to find cancer drivers, or genes, that, once mutated, give a selective advantage to a cancer cell, such as increased proliferation, suppression of apoptosis or the ability to avoid immune response. VHL is a key oncosuppressor gene for kidney cancer. Inactivation of the VHL gene is the most common event in renal carcinomas, accounting for 50–70% of sporadic cases (Scelo et al. 2014; Cancer Genome Atlas Research Network 2013; Thomas et al. 2006). Other drivers frequently mutated in kidney cancer include SETD2, a histone methylase which specifically trimethylated H3K4me3.

Tumor cells were shown to have altered DNA methylation profiles compared to normal cells of the same origins. Promoter DNA methylation can cause transcriptional inactivation of certain tumour suppressors, including the VHL gene in renal cell carcinomas. This event is mutually exclusive with VHL inactivation caused by deleterious mutations (Scelo et al. 2014; Cancer Genome Atlas Research Network 2013).

VHL gene is a key regulator of hypoxia. VHL is known to direct poly-ubiquitylation and further degradation of HIF1a, a transcription factor which activates hypoxia-associated genes (Maxwell et al. 1999; Tanimoto 2000). Therefore, inactivated or mutated VHL causes the accumulation of HIF1a transcription factor and subsequent activation of hypoxia transcription program. Interestingly, hypoxia itself is an important oncogenic factor: presence of hypoxia within a tumor is known to be associated with increased tumour malignancy, metastases rate and reduced overall survival (Jubb, Buffa, and Harris 2010).

DNA methylation is maintained and regulated by an oxidation-reduction reaction. The major DNA demethylation mechanism involves oxidation of methylcytosines to hydroxymethylcytosines with Tet enzymes which essentially are oxidoreductases that depend on Fe2+ and alpha-ketoglutarate (Ploumakis and Coleman 2015) and can be affected by an altered oxidation-reduction potential in hypoxia. It has recently been shown that DNA methylation is in fact affected under hypoxia (Thienpont et al. 2016). The levels of hydroxymethylated cytosine, an intermediate during cytosine demethylation, were drastically decreased in hypoxia. Interestingly, the CpGs which failed to be hydroxymethylated (and presumably demethylated afterwards) in hypoxic conditions were localized in promoters of important tumour suppressors thus preventing them from being activated. Reduced activity of hydroxymethylation process was also shown to cause an expected increase in overall DNA methylation at promoters.

In this work, we inactivated VHL gene in a renal carcinoma cell line Caki-1 with CRISPR/Cas9 editing to find out which consequences this common oncogenic event directly caused for gene expression and DNA methylation.

## Results

### VHL gene editing

We applied a CRISPR/Cas9 assay to introduce a homozygous frameshift deletion into VHL gene. This frameshift led to a premature stop in the C-terminal alpha domain of VHL. We checked how VHL editing affected the stability of Hif1a protein. In fact, Hif1a levels dramatically increased after VHL inactivation, the effect was much higher than that caused by hypoxia (Figure 1A).

**Figure 1.**
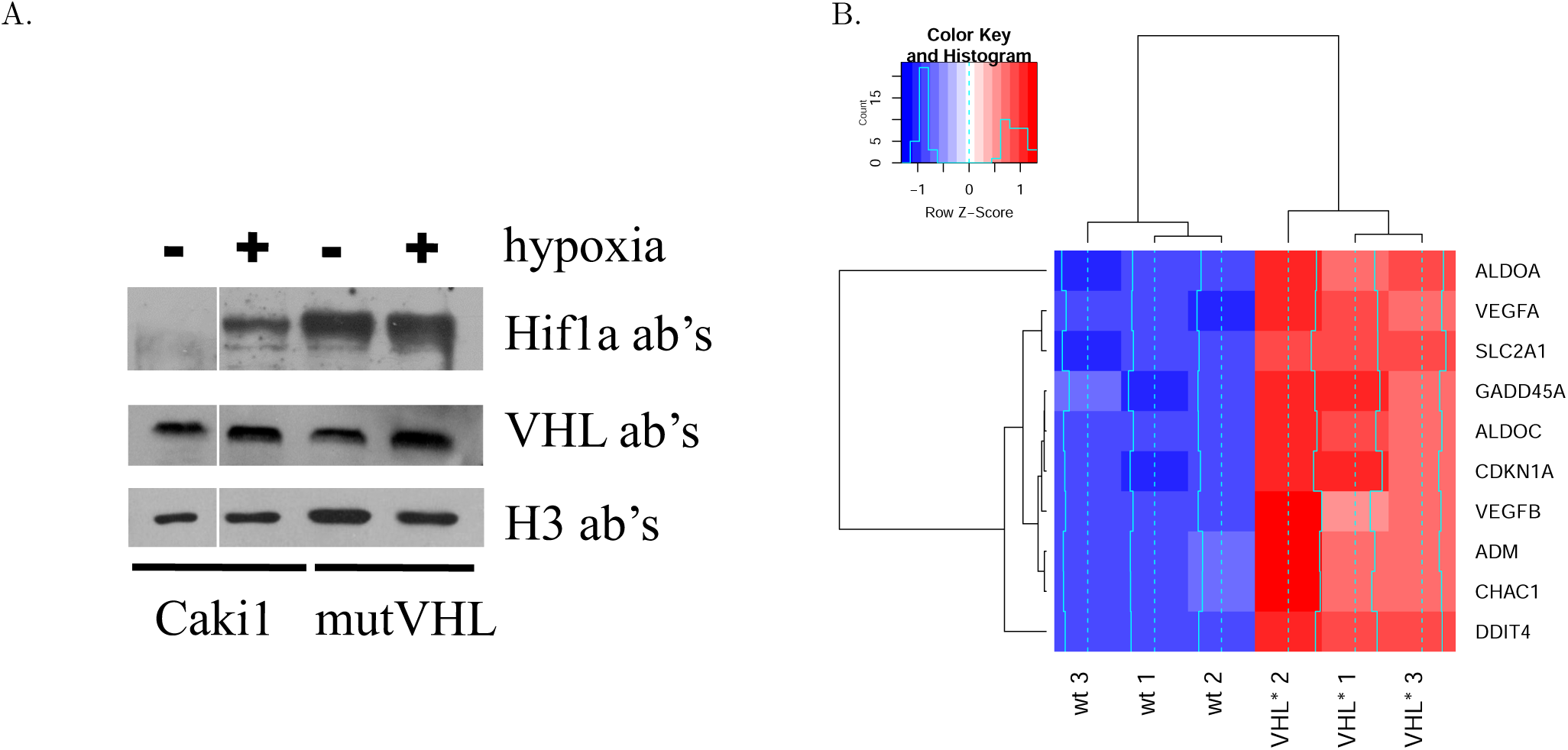
(A) Western blot depicting the level of Hif1a and Vhl protein in dependence of VHL mutation and hypoxia. (B) Heatmap showing the changes in expression of the hypoxia-associated genes. Hypoxia-inducible genes were activated in VHL* cells.

### VHL inactivation caused transcriptional upregulation of hypoxia-associated genes

We first checked if VHL inactivation led to a transcriptional response similar to that caused by hypoxia conditions. We took a conventional set of genes which were associated with hypoxia (Zhigalova et al. 2015; Eustace et al. 2013) and explored their expression in VHL wild type Caki-1 cells and VHL* cell lines after VHL editing. All of the considered genes showed significant upregulation (Figure 1B), which corresponded to our findings previously obtained by RT-qPCR for a different clone with VHL gene inactivated by editing (Zhigalova et. al, 2016. in press).

### VHL inactivation caused overall genome hypermethylation

We compared DNA methylation between VHL-wild type Caki-1 cell lines and the clone of Caki-1 cell line with VHL gene inactivated by a frameshift introduced by CRISPR/Cas9 gene editing (further referred as VHL*). We asked how DNA methylation of individual CpG dinucleotides was affected by VHL inactivation. Surprisingly, we observed a dramatic increase in genomic DNA methylation: most of the CpGs which significantly changes DNA methylation appeared to increase it (Figure 2).

**Figure 2.**
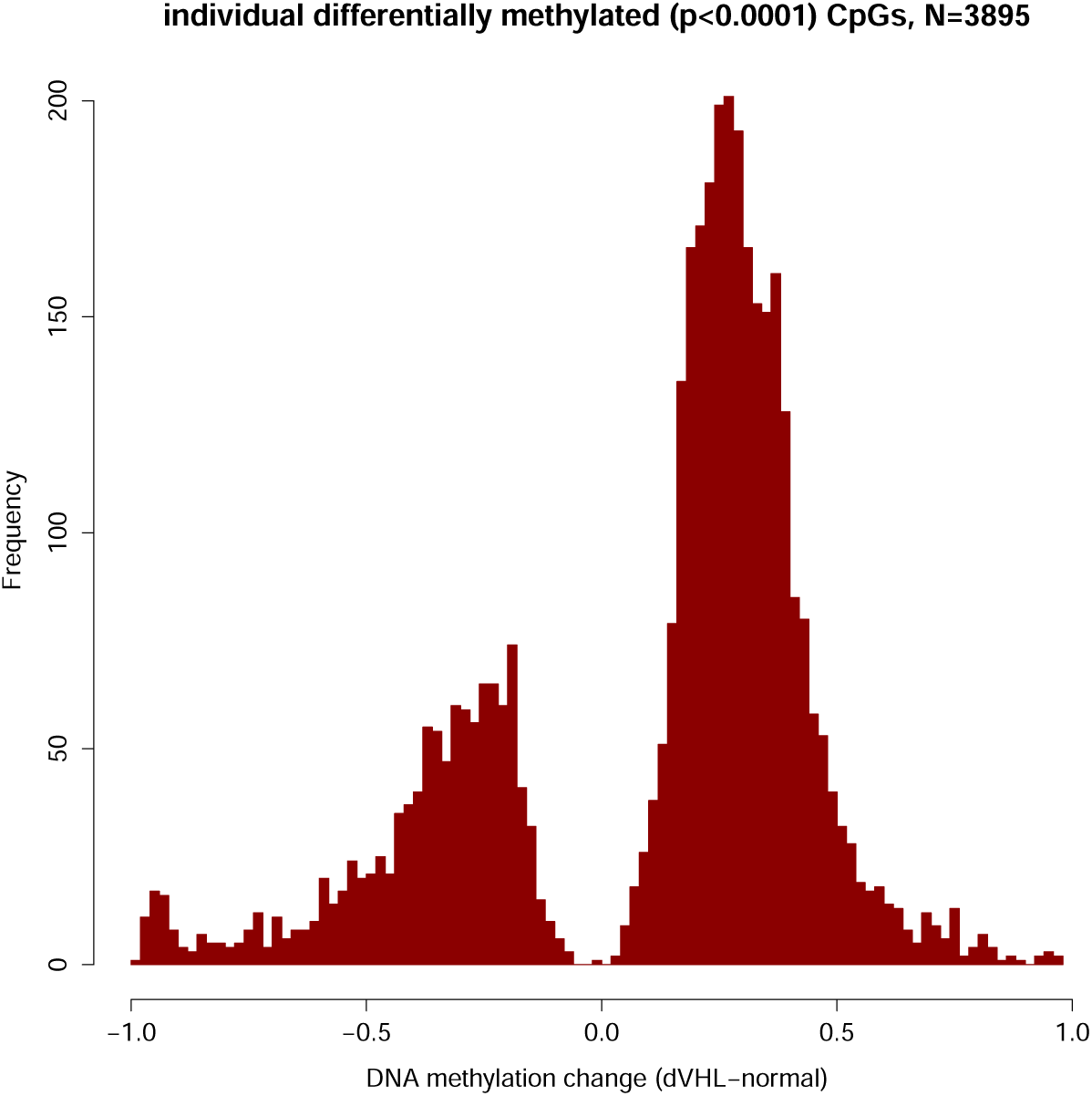
Distribution of changes in DNA methylation rates at individual CpGs following VHL inactivation. Only CpG positions which significantly changed their DNA methylation level were considered. DNA is on average hypermethylated after VHL inactivation.

### Hypermethylated DMRs were associated with chromatin modifiers

We discovered differentially methylated regions (DMRs) associated with VHL inactivation. Among 125 discovered DMRs, 86 significantly increased and 39 significantly increased DNA methylation in VHL* cells. The vast majority of the DMRs (81 out of 125 DMRs) was associated with transcription start sites (TSS) of known genes. Figure 3A shows what gene expression changes happen in the genes associated with a DMRs. Overall, a trend associating increased promoter methylation with decreased gene expression can be observed, though the effect is not significant due to several outliers. We studied which gene categories (GO) were enriched by the genes associated with the DMRs. Interestingly, the DMRs hypermethylated in VHL* cells were associated with genes encoding transcription factors (GO categories Transcription regulation, Transcription, Nucleotide binding). Moreover, we showed that expression of the six genes each falling in those categories was decreased after VHL inactivation (Figure 3B).

**Figure 3.**
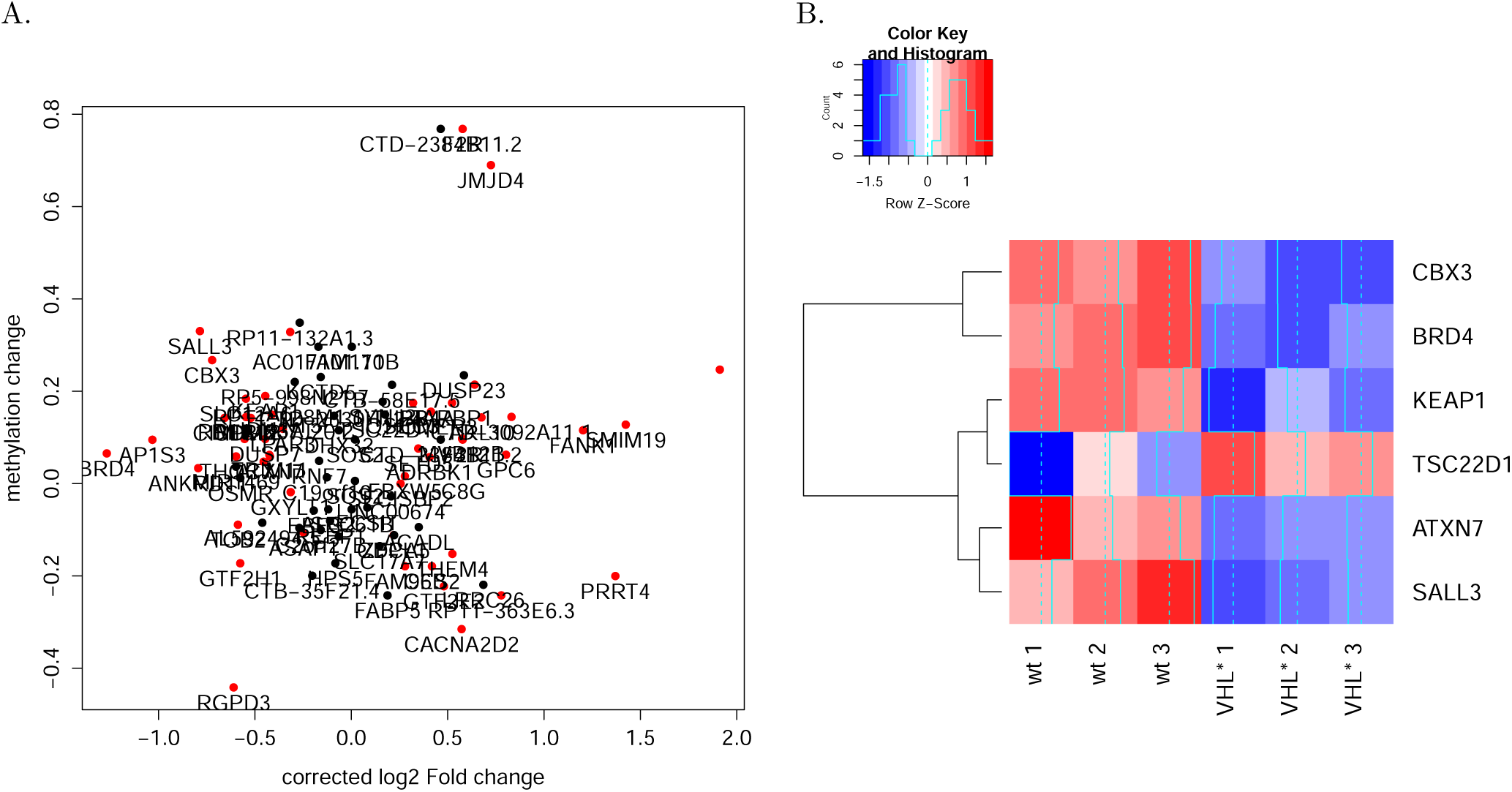
DMRs associated with gene TSSs. (A) Relation between gene expression fold-change and the change in DNA methylation within a DMR located near gene TSS. (B) Gene expression of transcription-related genes which were associated with a significantly hypermethylated DMR.

### AP-1 binding sites were enriched with hypermethylated CpGs

To attribute the observed DNA methylation changes to certain epigenetic mechanisms, we explored DNA methylation changes within different genomic and epigenomic markups. For each set of genomic intervals, we calculated a fraction of significantly hypo- or hypermethylated CpGs among all CpGs located within a given set of genomic intervals for which DNA methylation level had been profiled in the experiment. We started with genome segmentation performed by ChromHMM which subdivides genome into non-overlapping regions representing chromatin state (e.g., promoter or enhancer) according to the data on multiple epigenetic features. The highest fraction of hypermethylated CpGs was observed in inactive promoters (PromP) and weak or unconfirmed enhancer (DnaseU), while active enhancers show rather unchanged DNA methylation pattern with equal rate of hypo- and hypermethylation (Figure 4A).

**Figure 4.**
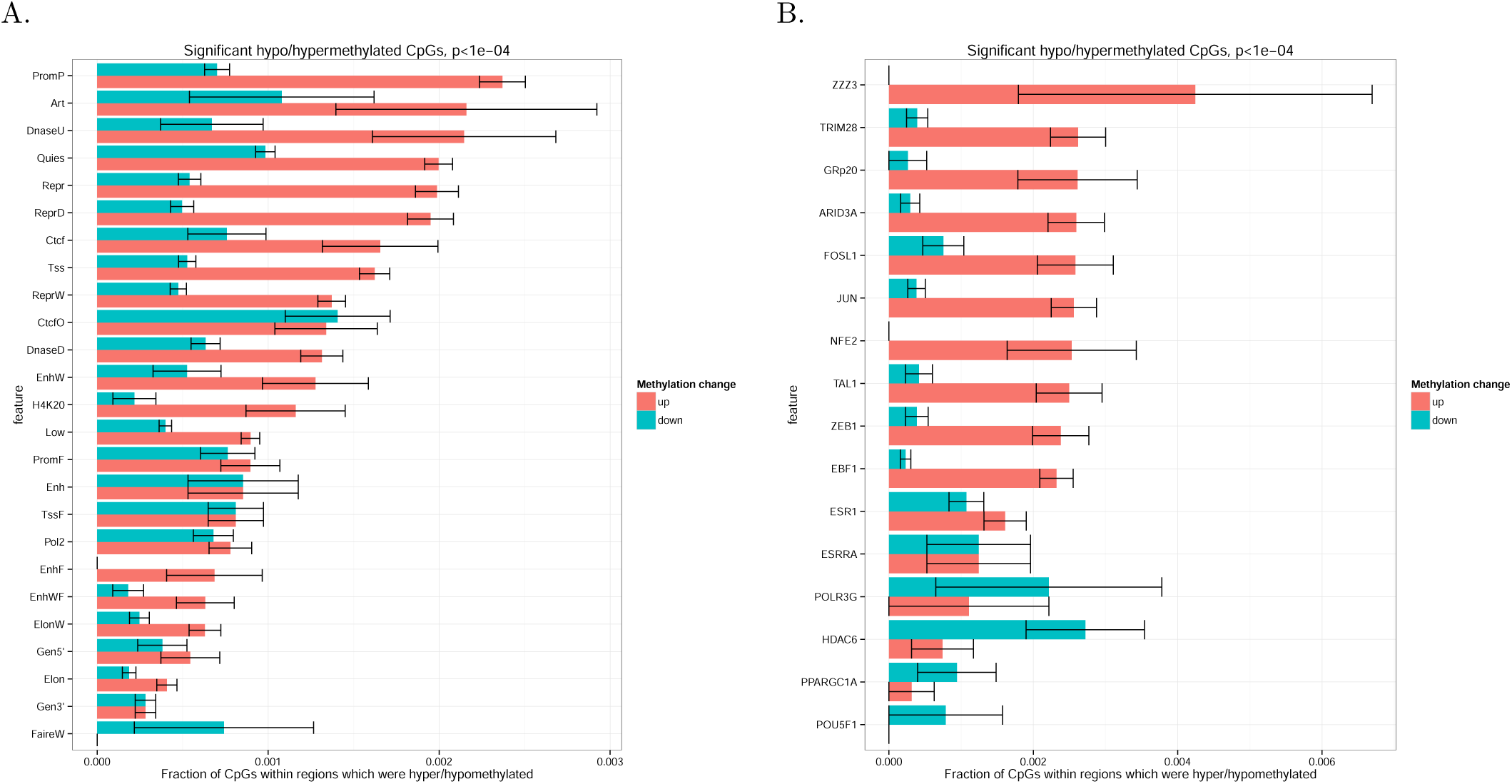
Distribution of hypo- and hypermethylated CpGs within certain epigenomic features. Fraction of significantly hypo- (blue bars) and hypermethylated (red bars) CpGs among all CpGs within a given set of genomic regions. (A) Regions of ChromHMM genome segmentation. (B). Transcription factor binding sites (only the sites with the highest hyper- and hypomethylation rates are plotted). CpGs that were significantly hypermethylated after VHL inactivation, were enriched in AP-1 (JUN/FOS) and TRIM28 binding sites. Hypomethylated CpGs were enriched in HDAC6 binding sites.

To understand if the overall genome hypermethylation could be caused by transcription factors, particularly by those involved in hypoxia response programme, we considered a comprehensive set of transcription factor binding sites annotated by ENCODE project (Figure 4B). We also included occurrences of HIF1a binding motif as HIF1a level was increased in VHL* cells. DNA methylation changes within HIF1a domains were not different from that observed genome-wide. Interestingly, the highest fraction of hypermethylated CpGs was observed in TRIM28 and AP-1 (JUN and FOS) binding sites, while only in HDAC6 binding sites we observed hypomethylation rather than hypermethylation.

To investigate how DNA methylation changes in the binding sites corresponded to gene expression changes of the respective transcriptional factors, we studied gene expression of FOS, JUN, TRIM28 and HDAC6 (Figure 5). All of the considered genes had significant expression changes according to DESeq2 test. Even though the genes showed different direction of expression changes, an apparent pattern could be observed: overexpression of activators (such as JUN and FOS genes forming together AP-1 transcription factor) and decreased expression of repressors (such as TRIM28) caused an increase in DNA methylation at their respective sites, while overexpression of a histone deacetylase HDAC6, involved in epigenetic repression, caused a decrease in DNA methylation at its sites. Therefore, the regions which could become more active due to an overexpression of a transcriptional activator or a decreased expression of transcriptional suppressor were preferred targets of DNA hypermethylation. This pattern can be explained by a hypothesis that DNA methylation in active chromatin is more prone to the processes causing DNA hypermethylation in VHL* cells.

**Figure 5.**
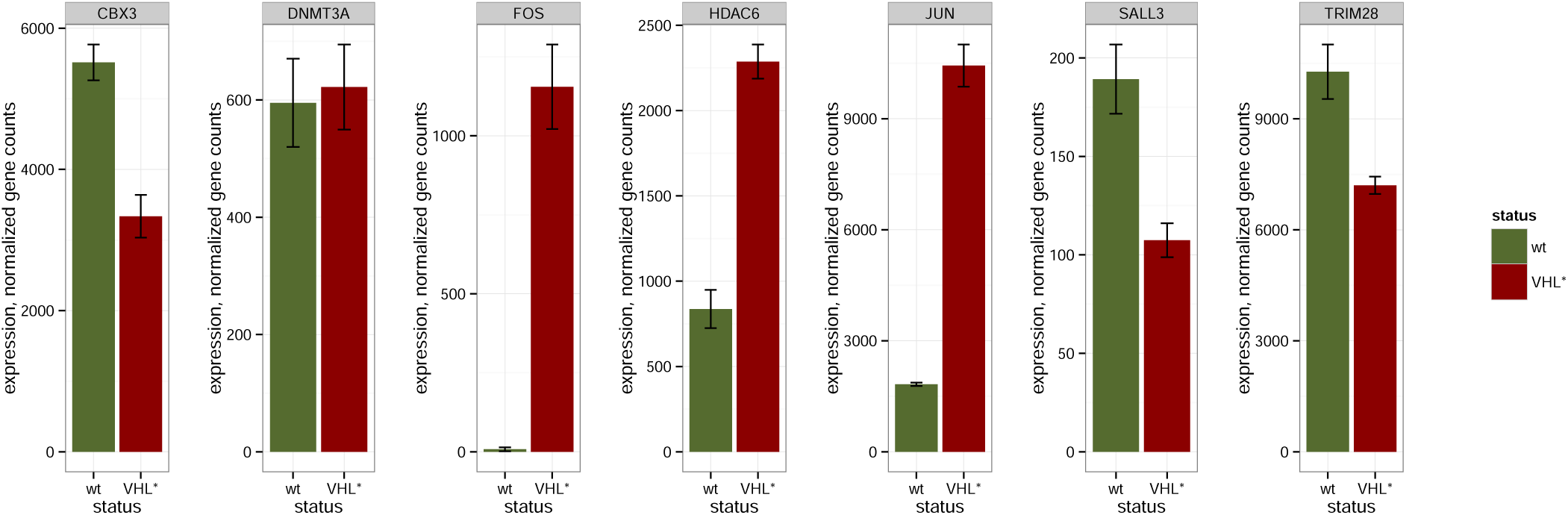
Gene expression in wild type Caki-1 and VHL* cells for certain genes either involved in regulation of DNA methylation or having their binding sites enriched with hypomethylated or hypermethylated CpGs.

## Discussion

It has recently been proposed that hypoxic conditions themselves can affect the functioning of the TET enzyme that is essentially an oxidoreductase (Thienpont et al. 2016). Low oxygen concentrations caused the decrease of hmC levels and subsequently led to overall hypermethylation at certain genomic loci. This effect was claimed to be independent on hypoxia signaling pathways, but rather was biochemically associated with decreased redox potential in hypoxia. Therefore, it was important to investigate if similar effect could be observed in conditions which are similar to hypoxia in terms of gene activation and cell signalling, but different due to the absence of actual hypoxia. VHL inactivation can be a model of such conditions and also represents an important oncogenic event.

The regions with significantly altered DNA methylation in VHL* cells were located at transcription start sites of the genes which were transcription factors and chromatin modifiers. The expression of these genes, apart of some outliers, is anticorrelated with DNA methylation. Moreover, some of the genes (like SALL3) were known to affect DNA methylation. This can arguably form feedback loops in which expression of a certain epigenetic modifiers is, on one hand, affected by DNA methylation and, on the other hand, influence genomic DNA methylation.

We initially assumed that increased abundance of HIF1a transcription factor would result in an altered DNA methylation pattern in its binding sites (Schödel et al. 2011; Tausendschön et al. 2015). Interestingly, HIF1a motif had a CG site which could be methylated. However, we didn't observe any changes in DNA methylation in HIF1a binding sites discovered by ChIP-seq. As the increased abundance of HIF1a could potentially result in emergence of novel HIF1a binding sites, we additionally checked what happened to DNA methylation in the occurrences of HIF1a motif in the genome. Overall methylation change in the motifs didn't differ from that observed genome-wide.

In an attempt to attribute the changes of DNA methylation to certain genomic and epigenetic features, we discovered that, among other transcription factor binding sites, the highest rate of hypermethylation was observed in AP-1 (Jun/Fos) binding sites. Surprisingly, Jun and Fos genes showed increased gene expression after VHL inactivation. It could be expected that this would lead to activation of their targets and subsequent decrease in methylation around their sites. However, an opposite effect was observed. To explain this contradiction, we formulated a hypothesis that DNA hypermethylation occurred genome-wide, but the regions of more open chromatin were more accessible for DNA methylase machinery and therefore they were more preferred targets of methylation. This hypothesis was even further confirmed by decreased expression of a transcriptional repressor TRIM28 and hypermethylation of its site. The opposite effect was detected for HDAC6 binding sites: they were the only epigenetic features which showed overall DNA demethylation rather than hypermethylation. This also supports the formulated hypothesis as increased expression of an epigenetic repressor HDAC6 coincided with hypomethylation of its sites.

## Methods

### CRISPR/Cas9-based gene editing

VHL gene editing with CRISPR/Cas9 was performed as described in our previous paper (Zhigalova et. al, 2016, in press). We introduced a homozygous frameshift mutation which led to a stop gain in the C-terminal alpha domain of VHL. Sanger sequencing confirmed that on the protein level, the original sequence VRSLYESGRPPKCAERPGAADTGAHCTSTDGRKLISVETYTVSSOLLMetVLMet-SLDLDTGLVPSLVSKCLILRVK(Stop) was substituted by VRSLYE(Stop).

### DNA methylation analysis

Three individual biological replicates from Caki-1 cell line and VHL* clone were taken for bisulfite sequencing. DNA methylation was profiled by reduced representation bisulfite sequencing (RRBS). Two micrograms of genomic DNA from a sample were digested using 60 U MspI (Fermentas, USA) in 50 *μ*l at 37°C for 18–24 h, followed by QIAquick purification (Qiagen, Germany). The end of the digested DNA was repaired, and an adenine was added to the 3' end of the DNA fragments according to the Illumina standard end repair and add_A protocol (Illumina, USA). Pre-annealed forked Illumina adaptors containing 5′- methylcytosine instead of cytosine were ligated to both ends of DNA fragments using standard Illumina adaptor ligation protocol (Illumina, USA). Ligated fragments were then separated by 2% agarose gel (Sigma-Aldrich, USA). Fragments between 170 bp and 350 bp, (includes adaptor length), were selected and cut from the gel. DNA from gel slices were purified using the Qiagen Gel Extraction Kit (Qiagen, Germany). The sodium bisulfite treatment and subsequent clean-up of size selected DNA was performed with the EZ DNA MethylationTM Kit (ZymoResearch, USA) according to the manufacturer's instructions. The bisulfite-treated DNA fragments were amplified using PCR and the following reaction: 5 *μ*l of eluted DNA, 1 of NEB PE PCR two primers (1.0 and 2.0) and 45 *μ*l Platinum PCR Supermix (Invitrogen, USA). The amplification conditions were as follows: 5 min at 95°C, 30 s at 98°C then 15 (10 s at 98°C, 30 s at 65°C, 30 s at 72°C), followed by 5 min at 72°C. The PCR reaction was purified by MinElute PCR Purification Kit (Qiagen), and final reduced representation bisulfite library was eluted in 15 *μ*l EB buffer. The concentration of the final library was measured using the Agilent 2100 Bioanalyzer (Agilent Technologies, USA). The library was sequenced on Illumina 2500 platform according to standard Illumina cluster generation and sequencing protocols. 100-bp single-end reads were generated. Reads were mapped to hg19 reference genome with Bismark software (Krueger and Andrews 2011).

Differential methylation analysis was performed by MethPipe/RADmeth software package (Song et al. 2013; Dolzhenko and Smith 2014). It is one of the tools for bisulfite-seq analysis which account for the fact that they are dealing with fractions of counts (e.g., for a particular CpG position, *k* reads support DNA methylation of the total of *N* reads covering the position). The tool operates with counts rather than with methylation rates 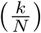 which are essentially prone to biases due to different coverage of a given position in each sample. It models the counts with beta-binomial distribution thus accounting for both binomially distributed shot noise in read counts and biological variance in DNA methylation of a given position between samples: *k* ~ *Binomial(N,p)*, where *p* ~ *Beta*(*α, β*), and *α* and *β* are fitted parameters. MethPipe/RADmeth was applied to bisulfite alignments as described in the manual. DNA methylation counts were merged between the forward and the reverse strand for each CpG thus excluding potentially mutated CpG sites. We first found differentially methylated individual CpG positions. Next, the positions were aggregated to differentially methylated regions (DMRs) with the default parameters.

ENSEMBL gene annotation (version 81) was used to find genes with TSS localized no further than 1 kilobase from differentially methylated regions.

### RNA-seq analysis

For RNA-seq, we used the same sample collection and treatment procedure as described for bisulfite sequencing. Four fish from each of the four experimental groups were taken for transcriptome analysis. Gills were isolated and fixed with IntactRNA^®^ reagent (Evrogen).

Total RNA was extracted from the samples with Trisol reagent according to the manufacturer's instructions (Invitrogen). Quality was checked with the BioAnalyser and RNA 6000 Nano Kit (Agilent). PolyA RNA was purified with Dynabeads^®^ mRNA Purification Kit (Ambion). An Illumina library was made from polyA RNA with NEBNext^®^ mRNA Library Prep Reagent Set (NEB) according to the manual. Paired-end sequencing was performed on HiSeq1500 with 2x75 bp read length. Approximately 25 million reads were generated for each sample.

Reads were mapped to hg19 genome with tophat2 software (version 2.1.0) (Kim et al. 2013). Gene models of non-overlapping exonic fragments were taken from ENSEMBL 54 database. For each exonic fragment, total coverage by mapped reads in each sample was calculated with bedtools multicov tool (version 2.17.0). Total gene coverage was calculated as a sum of coverages of all non-overlapping exonic fragments of a gene. Differential expression analysis was performed by applying default read count normalization (estimateSizeFactors) and performing per-gene negative binomial tests (nbinomTest), implemented in DESeq R package (version 1.22.0), with default parameters (Anders and Huber 2010). For each gene, the package provided both p-values and FDRs (p-values after Benjamini-Hochberg multiple testing corrections).

### Transcription factor motifs

PWM for HIF1a binding motif was downloaded from HOCOMOCO database (Kulakovskiy et al. 2012; Kulakovskiy et al. 2016). Occurrences of HIF1a binding motif were searched in hg19 genome with SARUS software (Kulakovskiy et al. 2010).

### Epigenomics tracks

Epigenomic tracks for hg19 genome were downloaded from ENCODE for HepG2 and GM12878 cell line. In particular, we studied peaks histone marks (wgEncodeBroadHistone tables), ChromHMM genome segmentation track for HepG2 cell line, and clustered transcription factor binding sites, conserved across many cell lines (wgEncodeRegTfbsClusteredV3)

## Supporting Information

**S1 Figure**

**Figure S1.**
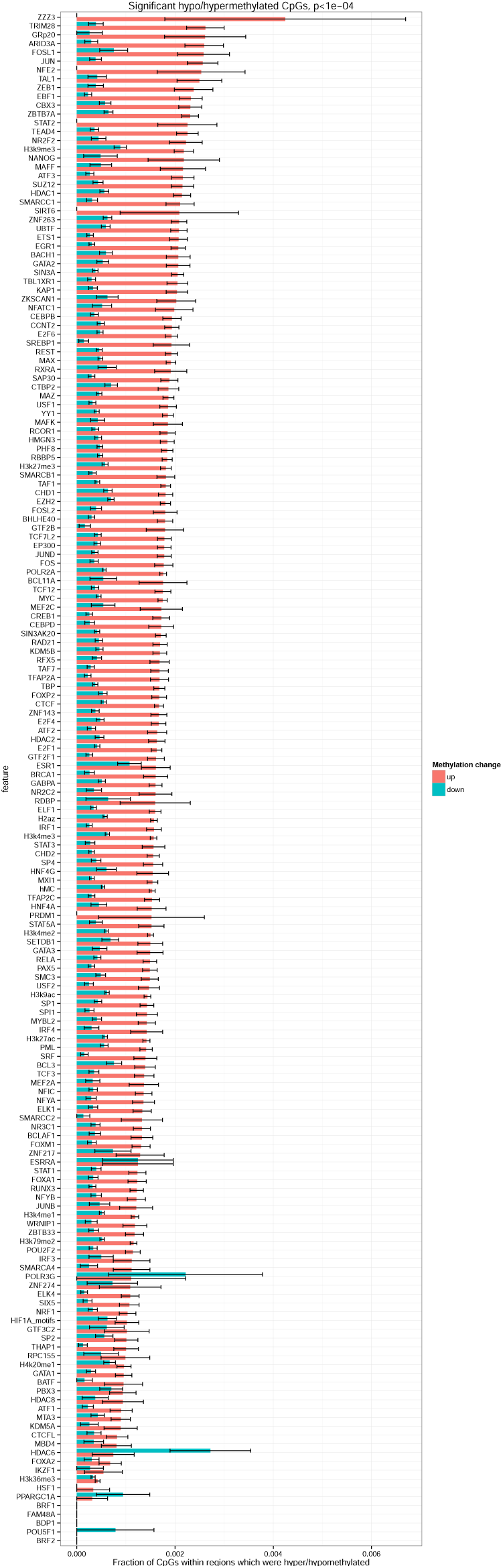
Distribution of hypo- and hypermethylated CpGs within certain epigenomic features. Fraction of significantly hypo- (blue bars) and hypermethylated (red bars) CpGs among all CpGs within a given set of genomic regions. (A) Regions of ChromHMM genome segmentation. (B). Transcription factor binding sites. CpGs that were significantly hypermethylated after VHL inactivation, were enriched in AP-1 (JUN/FOS) and TRIM28 binding sites. Hypomethylated CpGs were enriched in HDAC6 binding sites.

## References

Anders, Simon, and Wolfgang Huber. 2010. “Differential Expression Analysis for Sequence Count Data.” Genome Biology 11 (10): R106.

Cancer Genome Atlas Research Network. 2013. “Comprehensive Molecular Characterization of Clear Cell Renal Cell Carcinoma.” Nature 499 (7456): 43–49.

Dolzhenko, Egor, and Andrew D. Smith. 2014. “Using Beta-Binomial Regression for High-Precision Differential Methylation Analysis in Multifactor Whole-Genome Bisulfite Sequencing Experiments.” BMC Bioinformatics 15 (June): 215.

Eustace, A., N. Mani, P. N. Span, J. J. Irlam, J. Taylor, G. N. J. Betts, H. Denley, et al. 2013. “A 26-Gene Hypoxia Signature Predicts Benefit from Hypoxia-Modifying Therapy in Laryngeal Cancer but Not Bladder Cancer.” Clinical Cancer Research: An Official Journal of the American Association for Cancer Research 19 (17): 4879–88.

Jubb, Adrian M., Francesca M. Buffa, and Adrian L. Harris. 2010. “Assessment of Tumour Hypoxia for Prediction of Response to Therapy and Cancer Prognosis.” Journal of Cellular and Molecular Medicine 14 (1-2): 18–29.

Kim, Daehwan, Geo Pertea, Cole Trapnell, Harold Pimentel, Ryan Kelley, and Steven L. Salzberg. 2013. “TopHat2: Accurate Alignment of Transcriptomes in the Presence of Insertions, Deletions and Gene Fusions.” Genome Biology 14 (4): R36.

Krueger, F., and S. R. Andrews. 2011. “Bismark: A Flexible Aligner and Methylation Caller for Bisulfite-Seq Applications.” Bioinformatics 27 (11): 1571–72.

Kulakovskiy, Ivan V., Ilya E. Vorontsov, Ivan S. Yevshin, Anastasiia V. Soboleva, Artem S. Kasianov, Haitham Ashoor, Wail Ba-Alawi, et al. 2016. “HOCOMOCO: Expansion and Enhancement of the Collection of Transcription Factor Binding Sites Models.” Nucleic Acids Research 44 (D1): D116–25.

Kulakovskiy, I. V., V. A. Boeva, A. V. Favorov, and V. J. Makeev. 2010. “Deep and Wide Digging for Binding Motifs in ChIP-Seq Data.” Bioinformatics 26 (20): 2622–23.

Kulakovskiy, I. V., Y. A. Medvedeva, U. Schaefer, A. S. Kasianov, I. E. Vorontsov, V. B. Bajic, and V. J. Makeev. 2012. “HOCOMOCO: A Comprehensive Collection of Human Transcription Factor Binding Sites Models.” Nucleic Acids Research 41 (D1): D195–202.

Maxwell, P. H., M. S. Wiesener, G. W. Chang, S. C. Clifford, E. C. Vaux, M. E. Cockman, C. C. Wykoff, C. W. Pugh, E. R. Maher, and P. J. Ratcliffe. 1999. “The Tumour Suppressor Protein VHL Targets Hypoxia-Inducible Factors for Oxygen-Dependent Proteolysis.” Nature 399 (6733): 271–75.

Ploumakis, Athanasios, and Mathew L. Coleman. 2015. “OH, the Places You'll Go! Hydroxylation, Gene Expression, and Cancer.” Molecular Cell 58 (5): 729–41.

Scelo, Ghislaine, Yasser Riazalhosseini, Liliana Greger, Louis Letourneau, Mar Gonzàlez-Porta, Magdalena B. Wozniak, Mathieu Bourgey, et al. 2014. “Variation in Genomic Landscape of Clear Cell Renal Cell Carcinoma across Europe.” Nature Communications 5 (October): 5135.

Schödel, Johannes, Spyros Oikonomopoulos, Jiannis Ragoussis, Christopher W. Pugh, Peter J. Ratcliffe, and David R. Mole. 2011. “High-Resolution Genome-Wide Mapping of HIF-Binding Sites by ChIP-Seq.” Blood 117 (23): e207–17.

Song, Qiang, Benjamin Decato, Elizabeth E. Hong, Meng Zhou, Fang Fang, Jianghan Qu, Tyler Garvin, Michael Kessler, Jun Zhou, and Andrew D. Smith. 2013. “A Reference Methylome Database and Analysis Pipeline to Facilitate Integrative and Comparative Epigenomics.” PloS One 8 (12): 093310.

Tanimoto, K. 2000. “Mechanism of Regulation of the Hypoxia-Inducible Factor-1alpha by the von Hippel-Lindau Tumor Suppressor Protein.” The EMBO Journal 19 (16): 4298–4309.

Tausendschön, Michaela, Michael Rehli, Nathalie Dehne, Christian Schmidl, Claudia Döring, Martin-Leo Hansmann, and Bernhard Brüne. 2015. “Genome-Wide Identification of Hypoxia-Inducible Factor-1 and -2 Binding Sites in Hypoxic Human Macrophages Alternatively Activated by IL-10.” Biochimica et Biophysica Acta 1849 (1): 10–22.

Thienpont, Bernard, Jessica Steinbacher, Hui Zhao, Flora D'Anna, Anna Kuchnio, Athanasios Ploumakis, Bart Ghesquière, et al. 2016. “Tumour Hypoxia Causes DNA Hypermethylation by Reducing TET Activity.” Nature 537 (7618): 63–68.

Thomas, George V., Chris Tran, Ingo K. Mellinghoff, Derek S. Welsbie, Emily Chan, Barbara Fueger, Johannes Czernin, and Charles L. Sawyers. 2006. “Hypoxia-Inducible Factor Determines Sensitivity to Inhibitors of mTOR in Kidney Cancer.” Nature Medicine 12 (1): 122–27.

Zhigalova, Nadezhda, Artem Artemov, Alexander Mazur, and Egor Prokhortchouk. 2015. “Transcriptome Sequencing Revealed Differences in the Response of Renal Cancer Cells to Hypoxia and CoCl 2 Treatment.” F1000Research 4 (December): 1518.

